# Dual Enzyme Inhibition Strategy for Type 2 Diabetes: Targeting LDH-A and α-Amylase with Flavonoid-Acarbose Combinations

**DOI:** 10.1101/2024.11.09.622828

**Authors:** Adrija Bose, Suvroma Gupta

**Affiliations:** Department of Biotechnology, Haldia Institute of Technology, ICARE Complex, Hatiberia, Haldia, West Bengal-721657; Khejuri college, Baratala, Purba Medinipur, West Bengal

**Keywords:** Flavonoids, hyperglycemia, acarbose, amylase, Lactate-dehydrogenase-A

## Abstract

Diabetes mellitus is a complex and widespread disease affecting over 100 million people globally. It increases the risk of severe complications such as heart attack, neuropathy, and retinopathy. While various therapies aim to manage the disease, one effective approach involves reducing the activity of starch-degrading enzymes, such as α-amylase, to limit the amount of free glucose in the body. Another target is lactate dehydrogenase-A (LDH-A), an enzyme involved in gluconeogenesis through its role in generating pyruvate.

In recent years, flavonoids have gained significant attention for their potential to inhibit both α-amylase and LDH-A. This study investigates the flavonoids present in green and black tea for their ability to inhibit porcine pancreatic amylase (PPA) and LDH-A, comparing their efficacy to established anti-diabetic drugs such as Acarbose and Metformin using computational methods.

In vitro experiments demonstrated that Quercetin shows potent anti-LDHA-A activity with an IC_50_ of 4.161 µM. Quercetin inhibits PPA with an IC_50_ value of 20 µM, while eriodyctiol and myricetin exhibit IC_50_ values of 22 µM and 24 µM, respectively. Additionally, quercetin was found to synergistically enhance the inhibitory effects of the commonly used α-amylase inhibitor, Acarbose.

## 1. Introduction

Type 2 diabetes mellitus (T2DM) is a chronic metabolic disorder that currently affects approximately 8.8% of the global population, with epidemiological data suggesting an increasing prevalence across both developed and developing nations ^1^. It has emerged as one of the most significant global health challenges, and it is now classified as the 12th leading cause of mortality worldwide^2^.T2DM is characterized by insulin resistance, impaired insulin secretion, or both, resulting in chronic hyperglycemia^3^. The pathological mechanisms of the disease involve a progressive dysfunction in pancreatic β-cells, alongside insulin resistance in peripheral tissues, such as skeletal muscle, liver, and adipose tissue^4^. As the disease advances, the sustained high levels of blood glucose (hyperglycemia) contribute to a cascade of detrimental effects on various organ systems^4^. Prolonged hyperglycemia results in microvascular complications, such as diabetic retinopathy, nephropathy, and neuropathy, as well as macrovascular complications, including cardiovascular disease, which is a leading cause of morbidity and mortality in patients with T2DM^5^. The pathophysiology of these complications is driven by mechanisms such as oxidative stress, inflammation, and endothelial dysfunction, all of which exacerbate tissue damage and organ failure over time^6^. In addition to these physical health impacts, uncontrolled T2DM can lead to significant metabolic imbalances, including dyslipidemia and hypertension^7^. The multifactorial nature of T2DM underscores the importance of early diagnosis, effective management of blood glucose levels, and comprehensive treatment strategies that address both glycemic control and associated risk factors. With the upward trend in the incidence and prevalence of T2DM, it has become a major public health issue, demanding sustained global efforts in both prevention and management to mitigate its far-reaching consequences.

Among the six primary classes of antidiabetic agents commonly employed in the treatment of T2DM namely, insulin, sulfonylureas, biguanides, aldose reductase inhibitors, thiazolidinediones, insulin-like growth factors, and glucosidase inhibitors—alpha-glucosidase inhibitors occupy a prominent position^8, 9^. These inhibitors are regarded as better than other antidiabetic drugs due to their additional benefits, including their mild pharmacological profile, their ability to act without disrupting key biochemical pathways, and their localized mechanism of action within the intestine^10^. The therapeutic effect of alpha-glucosidase inhibitors is achieved through the inhibition of alpha-amylase, an enzyme responsible for the hydrolysis of complex carbohydrates into maltose, isomaltose, and oligomaltose. Acarbose, a non-hydrolyzable sugar analogue, serves as a potent inhibitor of alpha-amylase, though its use is associated with some adverse effects^11^.

Long-term administration of acarbose for the management of type 2 diabetes mellitus (T2DM) can result in various gastrointestinal complications, including diarrhea, flatulence, and abdominal pain^12^. Acarbose shifts a portion of the metabolic load of starch digestion to the colon, where bacterial fermentation of undigested starch produces gases, leading to symptoms such as bloating and discomfort. These adverse effects frequently contribute to the discontinuation of acarbose therapy ^13^. To address this issue, natural α-amylase inhibitors derived from sources such as wheat, black tea, green tea, and white beans have been explored for their potential to manage and control postprandial hyperglycemia^14-16^. Another widely used class of antidiabetic drugs is the biguanides. Biguanides primarily function by reducing the amount of glucose absorbed from food, inhibiting the conversion of fats and amino acids into glucose, and increasing glucose excretion by the kidneys^17^. Metformin, a commonly prescribed biguanide, is recognized for its efficacy as an antidiabetic agent. It has been shown to target glycerophosphate dehydrogenase, an enzyme that plays a role in regulating biochemical pathways involved in the development of T2DM. However, the use of biguanides, including metformin, is associated with gastrointestinal side effects such as diarrhoea, vomiting, and abdominal cramps^18^. Additionally, prolonged use of biguanides can lead to decreased absorption of vitamin B12^19^.

Several enzymes and hormones play critical roles in the regulation of blood glucose levels, with their dysregulation contributing to hyperglycemia. This paper focuses on the enzyme amylase, which has emerged as a key therapeutic target due to its presence in saliva and its primary function in catalyzing the breakdown of starches into simpler carbohydrates that are readily absorbed by the body. The subsequent absorption of glucose into the bloodstream, resulting in elevated plasma glucose levels, is a defining feature of diabetes. Therefore, inhibiting amylase activity represents a promising approach to controlling blood glucose levels in individuals with diabetes, as it could reduce the rate of glucose absorption and mitigate hyperglycemia^20^. In addition to α-amylase, previous studies have highlighted the role of lactate dehydrogenase-A (LDH-A) in the development of type 2 diabetes. Lactate dehydrogenase (LDH) is an enzyme found in nearly all living cells and is crucial for cellular respiration, specifically in the conversion of glucose into adenosine triphosphate (ATP), which is essential for various metabolic processes. LDH catalyzes the reversible conversion of pyruvate to lactate, playing a central role in anaerobic metabolism. Given its involvement in diverse metabolic pathways, LDH is often used as a biomarker for various physiological and pathological conditions ^21^. Research indicates that elevated LDH activity disrupts normal glucose metabolism, negatively impacting insulin secretion from the β-cells of the pancreatic islets of β-Langerhans^22^. Furthermore, several clinical studies have reported increased lactate levels in the plasma of patients with diabetes, suggesting a link between aberrant LDH activity and impaired glucose homeostasis^23^. Therefore, LDH-A may be considered a potential therapeutic target for the treatment of type 2 diabetes.

According to the recommendations of the American Cancer Society, the American Heart Association, and the American Diabetes Association, an increased intake of plant-derived foods is a preventive measure against several chronic diseases, including type 2 diabetes, cancer, and cardiovascular complications^24^. Flavonoids, a class of secondary metabolites of plant origin, are widely found in fruits, vegetables, grains, bark, roots, stems, flowers, tea, and wine. These plant-derived polyphenolic compounds exhibit a broad range of biological activities, including free radical scavenging, antimicrobial, anti-inflammatory, antitumor, and anti-lipid peroxidation effects^25^. Interest in flavonoids has grown significantly following the discovery of the “French Paradox,” which refers to the lower mortality rates from cardiovascular disease observed in Mediterranean populations, despite a diet high in saturated fats. This phenomenon has been linked to the consumption of red wine, which contains a flavonoid known as quercetin^26^. Following this observation, extensive research has been conducted to isolate, identify, and understand the mechanistic pathways through which flavonoids exert their protective effects against various chronic diseases.

In addition to wine, tea is another flavonoid-rich beverage that is gaining attention for its potential health benefits. Tea is one of the most widely consumed beverages globally and is derived from the leaves of *Camellia sinensis*. These leaves are abundant in polyphenolic compounds, including flavonoids, catechins, and epigallocatechin. There are primarily two types of tea available commercially: black tea and green tea. The various flavonoids present in tea are illustrated in Fig. 2.

**Figure 1:**
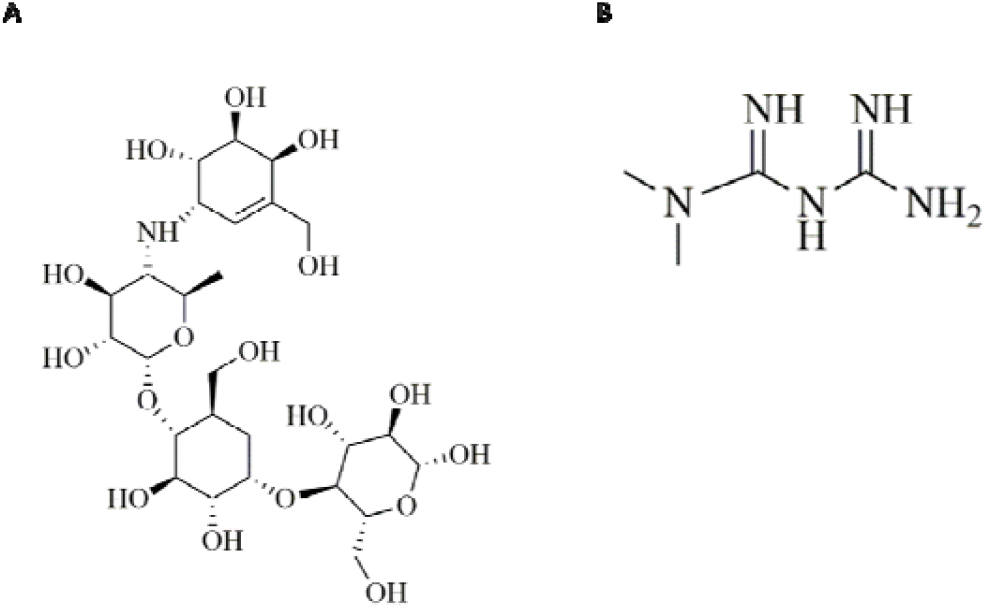
Chemical structures of (A) Acarbose and (B) Metformin

**Figure 2:**
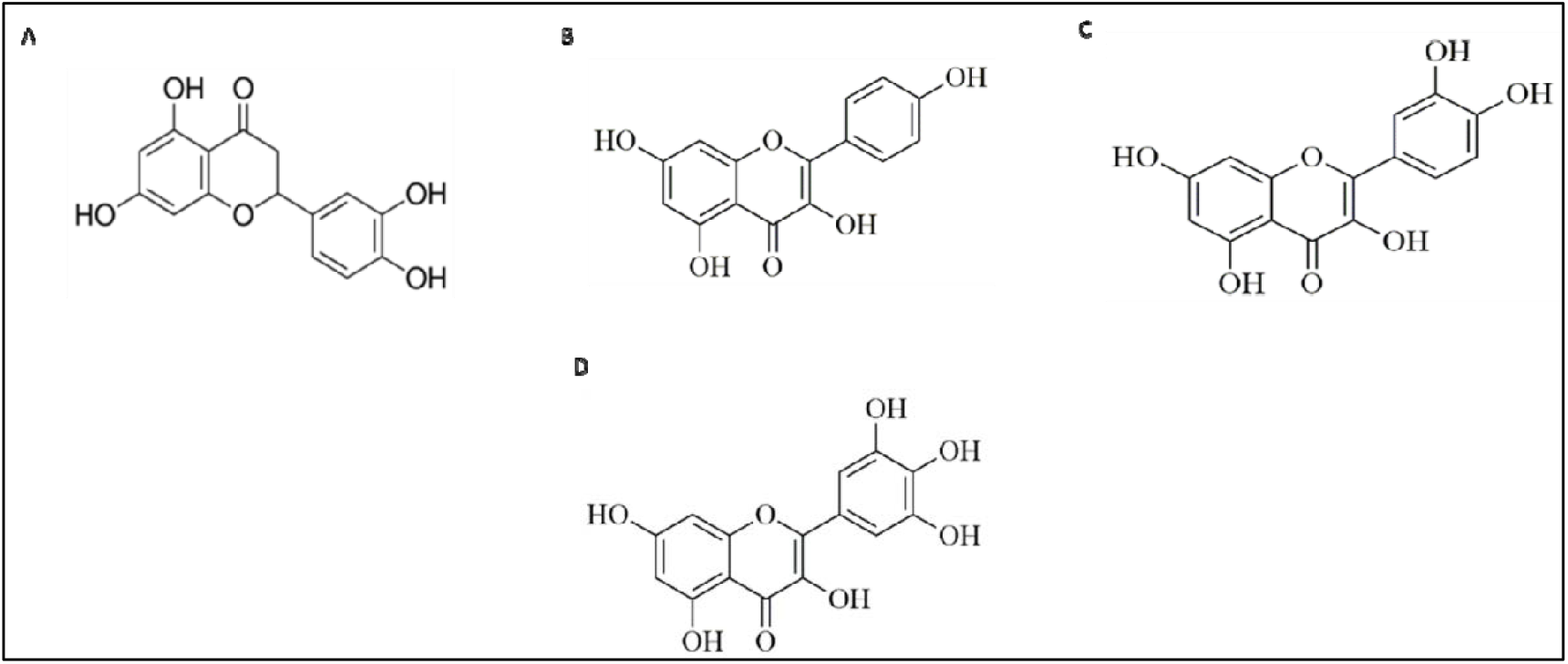
Chemical structures of (A) Eriodyctiol (B) Kaempferol (C) Quercetin (D) Myricetin

The present study was undertaken to evaluate the potential of various flavonoids in inhibiting α-amylase and lactate dehydrogenase-A (LDH-A) activities. The experimental work includes assessing the inhibitory effect of tea, considered as a single entity, on LDH-A activity. An *in silico* analysis was conducted to identify the most abundant flavonoid in tea with the highest binding affinity for LDH-A, followed by an inhibition assay to validate this finding. Furthermore, in silico experiments were performed to compare the binding energies of flavonoids with α-amylase, assessing their potential as anti-diabetic agents relative to established models, such as Acarbose and Metformin. Additional investigations were conducted to elucidate the kinetics of α-amylase inhibition by flavonoids and to estimate their synergistic inhibitory effects in combination with Acarbose.

## 2. Materials and Method

### 2.1. LDH-A Assay

The assay mixture contained 50mM Tris buffer (pH-7.6), 1.8mM NaPy, and 5mM NADH. A 1:10 diluted enzyme was used and the OD was measured at 340nm. The reaction was monitored from the decrease in the OD 340 value for a period of time. The molar coefficient of NADH is assumed to be 6220 M^-1^cm^-1^ as per literature and the change in absorbance for the disappearance of 1 micromole of NADH was taken as 2.0733. Thus, the activity of the enzyme, per ml of CE was calculated as:

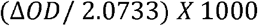

where Δ *OD* = *ODi* – *ODf* (3 min assay)

### 2.2. Molecular Docking

The target proteins for this study were α-amylase (PDB ID: 2QMK) and lactate dehydrogenase-A (LDH-A; PDB ID: 1D-5ZJF). Prior to docking simulations, the protein structures were prepared by removing water molecules and native ligands using AutoDock Tools (version 1.5.6)) ^27^. This step, referred to as protein cleaning or purification, ensures that extraneous molecules, which may interfere with ligand binding, are eliminated. Subsequently, partial charges and polar hydrogen atoms were added to the proteins to complete the preparation. The 3D conformations of the ligand molecules were retrieved from the PubChem database. Ligand preparation was performed using AutoDock Tools (version 1.5.6)) ^27^, during which Gasteiger charges were assigned to the ligands, and non-polar hydrogen atoms were merged. Docking simulations were conducted using AutoDock Vina) ^27^, employing its scoring function to evaluate binding interactions. The results were reported as binding scores to quantify docking accuracy. The visualization of protein-ligand interactions was performed using PyMOL (version 4.6.0)^28^ and Discovery Studio 2021^29^. Discovery Studio was further utilized to analyze the detailed interactions between the enzymes and the ligands at the binding sites.

### 2.3. Amylase Assay

Porcine pancreatic α-amylase and 3,5-dinitrosalicylic acid were purchased from Sigma Chemical Co. Ltd. (St. Louis, MO, USA). Acarbose was procured from Bayer (Leverkusen, Germany). Eriodictyol, Quercetin, and Myricetin had been purchased from Sigma. Na-K tartrate, potassium dihydro-orthophosphate, potassium hydroxide, sodium carbonate, sodium chloride, sodium dihydro-orthophosphate, sodium hydroxide) and dimethyl sulfoxide used was of analytical grade.

Porcine pancreatic α-amylase was dissolved in 50 mM phosphate buffer saline, pH 6.9. The various concentrations of flavonoids dissolved in DMSO were added to the assay mixture containing 1 g/L starch in 50 mM phosphate buffer saline. The reaction was initiated by adding PPA (3 U/mL) to the incubation medium to a final volume of 500 μL. The reaction was stopped after 10 min by adding 0.5 mL dinitrosalicylic (DNS) reagent (1% 3, 5-dinitrosalicylic acid, 0.2% phenol, 0.05% Na2SO3, and 1% NaOH) to the reaction mixture. The mixtures were heated at 100 °C for 10 min. After cooling to room temperature, absorbance (Abs) was recorded at 540 nm using a spectrophotometer. α-Amylase inhibition assays are conducted according to the following calculation.

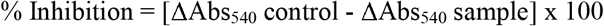

The extent of PPA inhibition had been calculated as IC_50_ values (the concentration of the flavonoid required to produce 50% inhibition of the test samples).

### 2.4. Determination of the PPA inhibitory mode of action of flavonoids

The kinetic model of inhibition of flavonoids against PPA had been estimated with a series of samples in which the concentration of the substrate was varied in the absence or presence of different concentrations of the inhibitors. The mode of inhibition (i.e., competitive, non-competitive, and uncompetitive) of the test flavonoids was determined from the nature of the curve using the Lineweaver-Burk plot. Respective K_m_ (dissociation constant) and V_max_ (maximum reaction velocity) values for each of the PPA assays were estimated from the slope and intercept of the curve ^30^. by plotting the inverse of PPA reaction velocity (V) versus 1/ [substrate (starch) concentration]

### 2.5. Effect of quercetin on in vitro PPA inhibition in combination with acarbose

To evaluate the effect of flavonoids on inhibiting PPA activity, quercetin was tested at its half-maximal inhibitory concentration (IC_50_), along with acarbose at reduced concentrations of 3 µM and 6 µM. The percentage of PPA inhibition was determined using the standard DNS assay, as previously described. The inhibitory activity of PPA was determined using the DNS (3,5-dinitrosalicylic acid) assay. In brief, the reaction mixture containing PPA enzyme, substrate (starch), and the test compounds (quercetin and acarbose) was incubated at 37°C for 30 minutes. After incubation, the reaction was terminated by adding DNS reagent, followed by boiling the mixture for 5 minutes to allow color development. The absorbance was measured at 540 nm, and the percentage inhibition of PPA was calculated by comparing the absorbance of the test samples to the control (without inhibitor).

### 2.6. In vitro anti-lipid peroxidation assay in the presence of flavonoids

Lipid peroxide formation has been measured using the modified method of Onkawa et al ^31^. Freshly excised goat liver has been homogenized in PBS (pH 6.9). 1 ml of liver homogenate (10%, w/v) had been added to the test samples containing flavonoids. Lipid peroxidation had been initiated after the addition of 100 µl of 15 mM FeSO_4_ followed by an incubation of 30 minutes at room temperature. 100 µl of the reaction mixture with test flavonoid was taken in a tube containing 1.5 ml of 10% TCA. After 10 minutes of centrifugation, the supernatant was mixed with 1.5 ml of 0.67 % TBA in 50% acetic acid. Following 30 minutes of boiling the intensity of the pink color, the solution had been measured at 535 nm. % of inhibition by flavonoids had been calculated as stated above.

### 2.7. Preparation of enzyme extraction from goat liver

Extraction buffer:10 ml (50mM) Tris HCl (pH 7.6), 1mM EDTA, and 1 mM β-mercaptoethanol were used and the final volume was adjusted up to 50 ml. The goat liver was homogenized in a homogenizer and centrifuged at 12000 rpm for 10 minutes at 4°C. The supernatant was collected discarding the pellet and the crude enzyme thus obtained was diluted 10 times for every use.

### 2.8. Binding Thermodynamics of Quercetin with PPA

Isothermal titration calorimetry was performed using a VP-ITC Microcalorimeter from MicroCal, Inc. located in Northampton, MA. PPA was extensively dialyzed using phosphate buffer, and the ligand Quercetin was dissolved in the final dialysate. The pH levels of the amylase protein and the ligand solution were adjusted to be identical before being loaded into the calorimeter. The heat generated solely from the dilution of the ligand in the buffer was removed from the titration data. The data was analyzed to determine the binding stoichiometry (N), affinity constant (Ka), and thermodynamic parameters of the reaction, utilizing Origin 5.0 software.

## 3. Results and Discussion

### 3.1. Inhibition of LDH-A by tea extracts

In a comparative analysis of green and black tea for their inhibitory effects on LDH-A activity, green tea demonstrated a more pronounced inhibitory effect within the tested concentration range of 0.25% to 0.5%. Although the inhibition exhibited by both teas was modest at lower concentrations, green tea consistently showed greater potency than black tea, particularly at the highest concentration tested (0.5%). Notably, green tea achieved nearly 80% inhibition of LDH-A activity at a concentration of 0.5%. In contrast, black tea, even at a higher concentration of 2%, exhibited significantly lower inhibitory potential as depicted in Fig 3 (A). This suggests that the flavonoid profile of green tea is likely more enriched in compounds with strong LDH-A inhibitory properties. The greater efficacy of green tea can be attributed to its higher content of catechins, such as epigallocatechin gallate (EGCG), which have been reported to exhibit potent enzyme inhibition, including against lactate dehydrogenase isoforms^17^. The observed difference in inhibition could also be influenced by the degree of oxidation that occurs during the processing of tea leaves. Green tea undergoes minimal oxidation, preserving the integrity of its flavonoid compounds, while black tea is fully oxidized^32^, which may result in the degradation or modification of certain bioactive compounds, reducing its inhibitory efficacy. These findings are consistent with previous studies that have reported superior antioxidant and enzyme-inhibitory activities of green tea compared to black tea ^33^.

**Figure 3:**
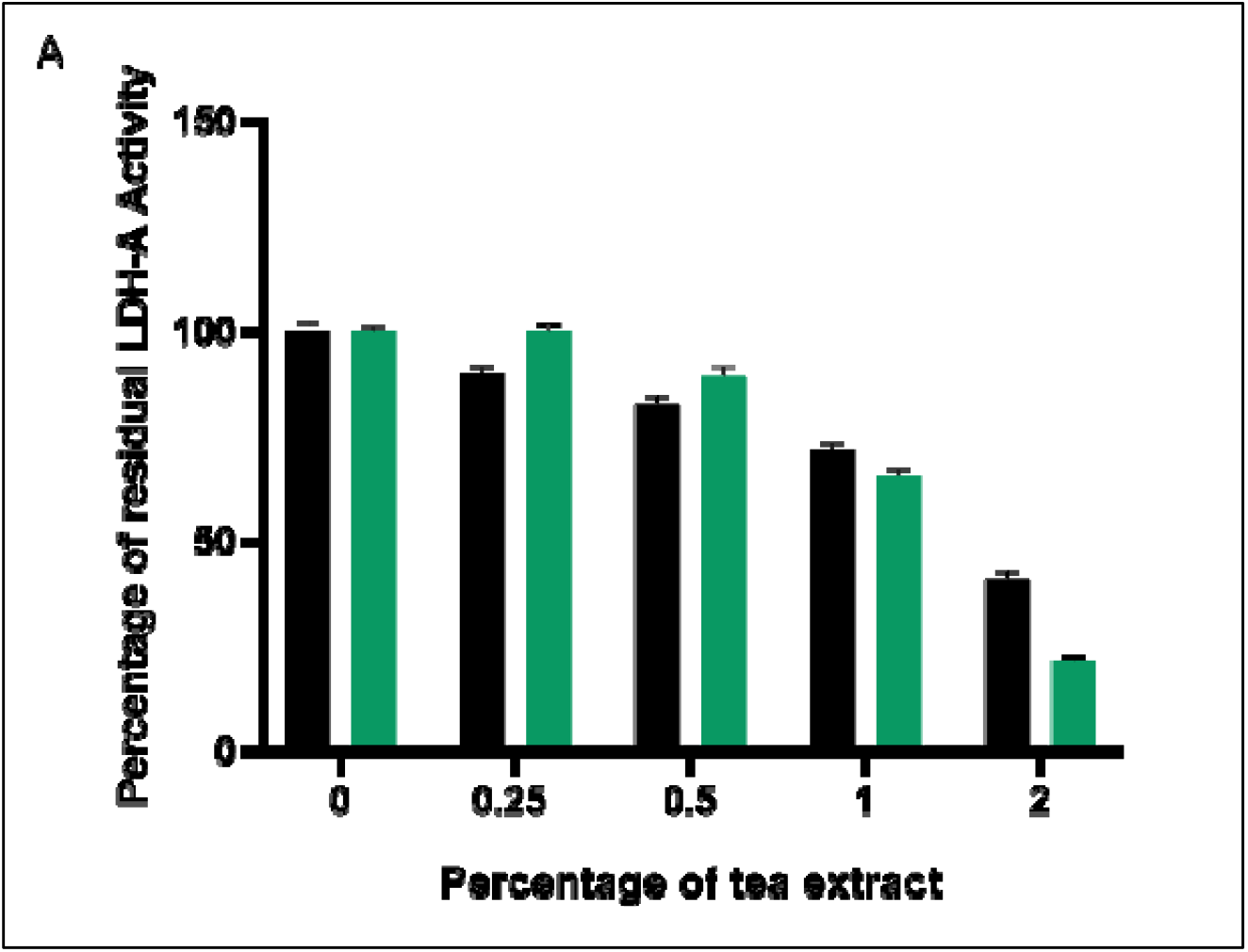
**A:** Effect of tea extract on LDH-A activity

### 3.2. Interactions of ligands with LDH-A

The analysis of protein-ligand interactions reveals diverse bonding patterns for the tested compounds, with varying degrees of stability conferred through hydrogen bonding, van der Waals interactions, and other non-covalent forces. Acarbose predominantly forms hydrogen bonds with Lysine and Glutamine, indicating stable interactions between the ligand and the protein. The formation of hydrogen bonds generally suggests favorable interactions that contribute to the stability of the protein-ligand complex. Additionally, Acarbose exhibits van der Waals interactions with Proline, Lysine, and Glutamine residues. However, it also shows an unfavorable acceptor-acceptor interaction with Alanine, which may slightly reduce the overall stability of the interaction. In comparison, Kaempferol displays a broader range of interactions. It forms hydrogen bonds with the polar amino acid residue Threonine and engages in van der Waals interactions with Valine, Threonine, Glutamine, Aspartate, Phenylalanine, Proline, and Glycine residues. Notably, Kaempferol also forms a π-alkyl bond with Valine and a π-π stacked interaction with Tryptophan, a cyclic amino acid. These interactions indicate a stable binding, although the predominant van der Waals interactions suggest weaker individual bonding forces compared to hydrogen bonds, but still contribute significantly to ligand-protein stability. Metformin, a widely used anti-diabetic drug, interacts with the protein primarily via conventional hydrogen bonds with Asparagine and Threonine, while additional stability is provided by van der Waals interactions with Tyrosine, Phenylalanine, Threonine, Tryptophan, Leucine, and Lysine. The combination of hydrogen bonding and van der Waals forces in Metformin’s interactions suggests a balanced and stable binding to the target protein. Myricetin, another flavonoid, shows extensive interactions with the protein, forming seven hydrogen bonds with polar residues such as Serine, Tyrosine, Leucine, and Alanine. These multiple hydrogen bonds significantly enhance the stability of the protein-ligand complex. Additionally, van der Waals interactions with Glycine, Leucine, Phenylalanine, Glutamine, Lysine, and Proline contribute further to its binding stability. Myricetin also exhibits three π-alkyl bonds, an amide-π stacked bond, and π-cation interactions with Lysine, suggesting a robust multi-interaction profile that enhances its inhibitory potential. Quercetin, another potent flavonoid, displays several stabilizing interactions with lactate dehydrogenase (LDH). A hydrogen bond is formed with Tyrosine, contributing to the stability of the protein-ligand structure. Additionally, π-alkyl bond formation is observed with Lysine and Leucine residues. Van der Waals interactions with Phenylalanine, Tyrosine, Threonine, and Tryptophan further contribute to the overall interaction profile. Quercetin also engages in π-sigma, π-π T-shaped, and π-π stacked bonds with Leucine, Tryptophan, and Tyrosine, respectively, indicating a highly stable and diverse set of interactions. Eriodyctiol demonstrates a mix of π-π T-shaped interactions, hydrogen bonds, van der Waals interactions, and Pi-alkyl interactions. However, Eriodyctiol also shows unfavorable donor-donor interactions, which could negatively affect the stability of the protein-ligand complex. The variety of interactions across these ligands suggests that flavonoids such as Myricetin and Quercetin show high potential for strong and stable binding with LDH, largely due to their extensive hydrogen bonding and additional stabilizing non-covalent interactions. This comparative analysis highlights the importance of diverse interaction profiles, particularly the role of hydrogen bonds and van der Waals forces in stabilizing protein-ligand complexes. A graphical representation of the protein-ligand interactions for each compound is provided in Fig. 4 (A-G).

**Figure 4:**
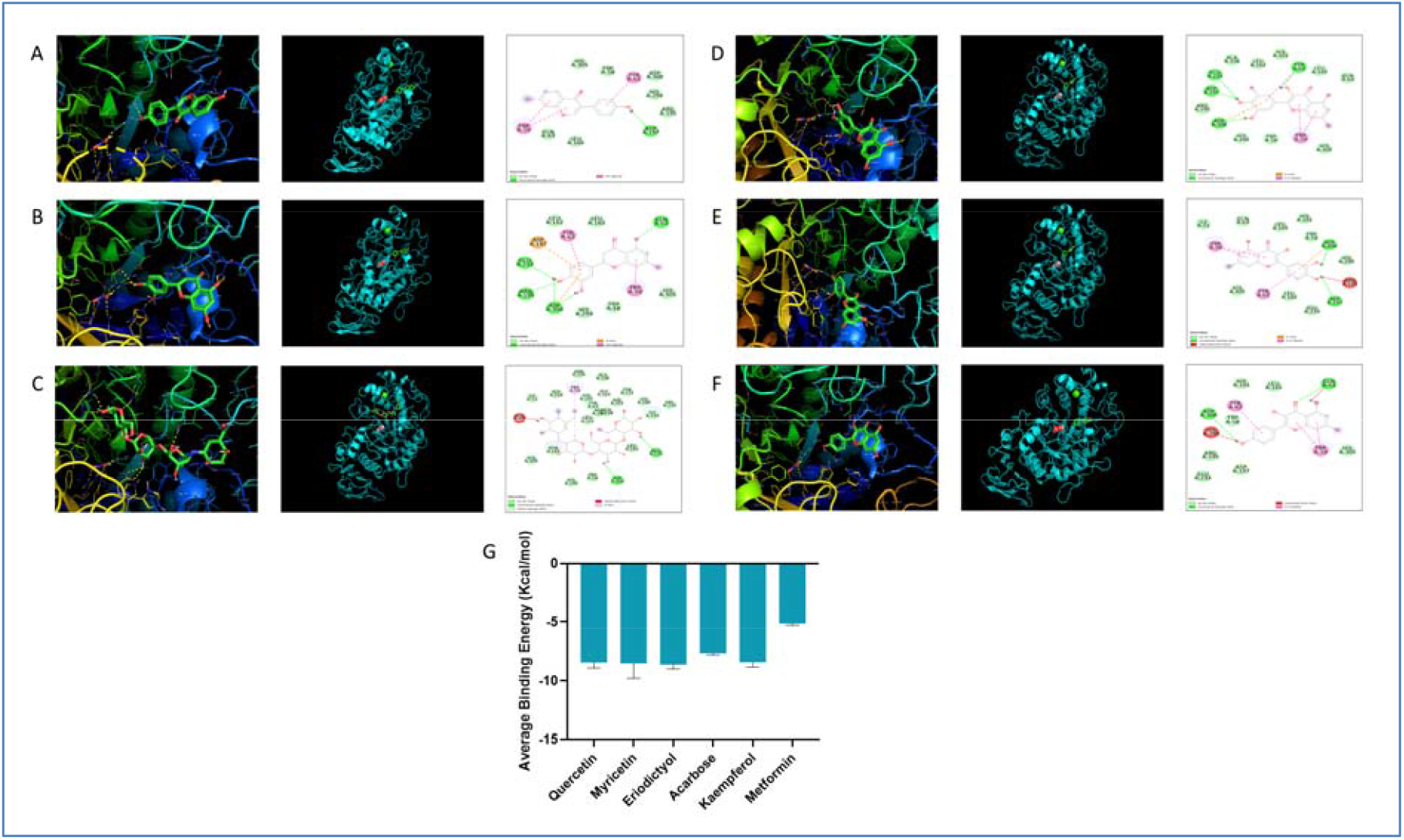
Computational visualization of the interaction of (A) Eriodyctiol (B) Acarbose (C) Kaempferol (D) Metformin (E) Myricetin and (F) Quercetin. (G) Binding affinity of selected ligands with LDH-A

### 3.3. Effect of Quercetin on LDH-A

Quercetin exhibited a dose-dependent inhibition of lactate dehydrogenase (LDH) activity, with a significant reduction in enzymatic activity observed as the concentration of quercetin increased. The result in Fig. 5 (A) suggests that a concentration of 10 µg, of quercetin inhibited approximately 40% of LDH activity compared to the control, demonstrating its potency as an LDH inhibitor. This reduction in activity suggests that quercetin has the potential to interfere with the enzyme’s function, potentially through direct binding to the active site or through allosteric modulation. Further analysis revealed that quercetin recorded an IC_50_ value of 4.161 µM. The mechanism of inhibition may be attributed to the ability of quercetin to form stabilizing interactions with key residues within the LDH active site, as evidenced by previous docking studies. Quercetin’s interactions, such as hydrogen bonding and π-π stacking with residues like Tyrosine and Tryptophan, likely play a critical role in its ability to disrupt LDH activity.

**Figure 5:**
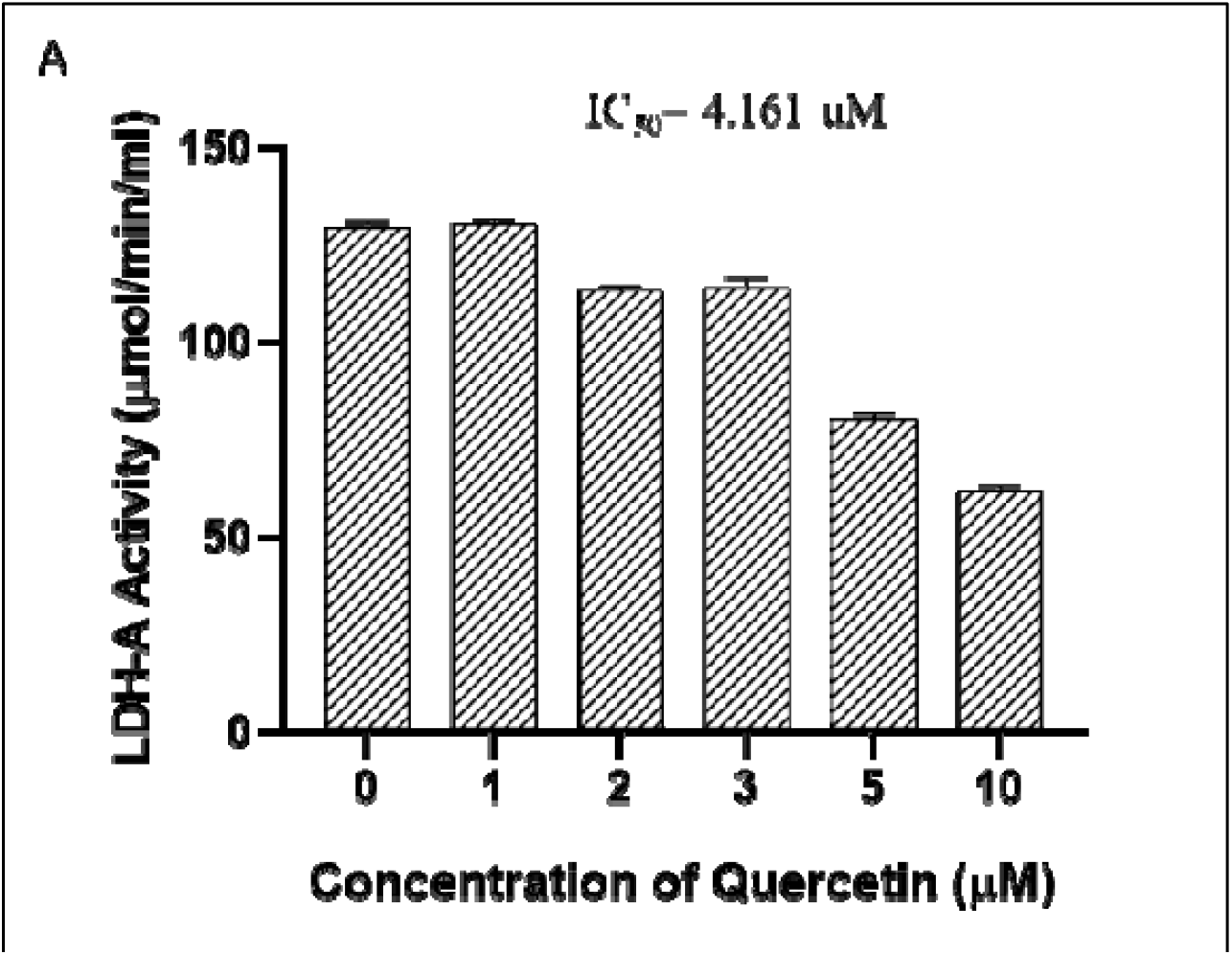
**A**: Inhibition of LDH-A by Quercetin

**Figure 6:**
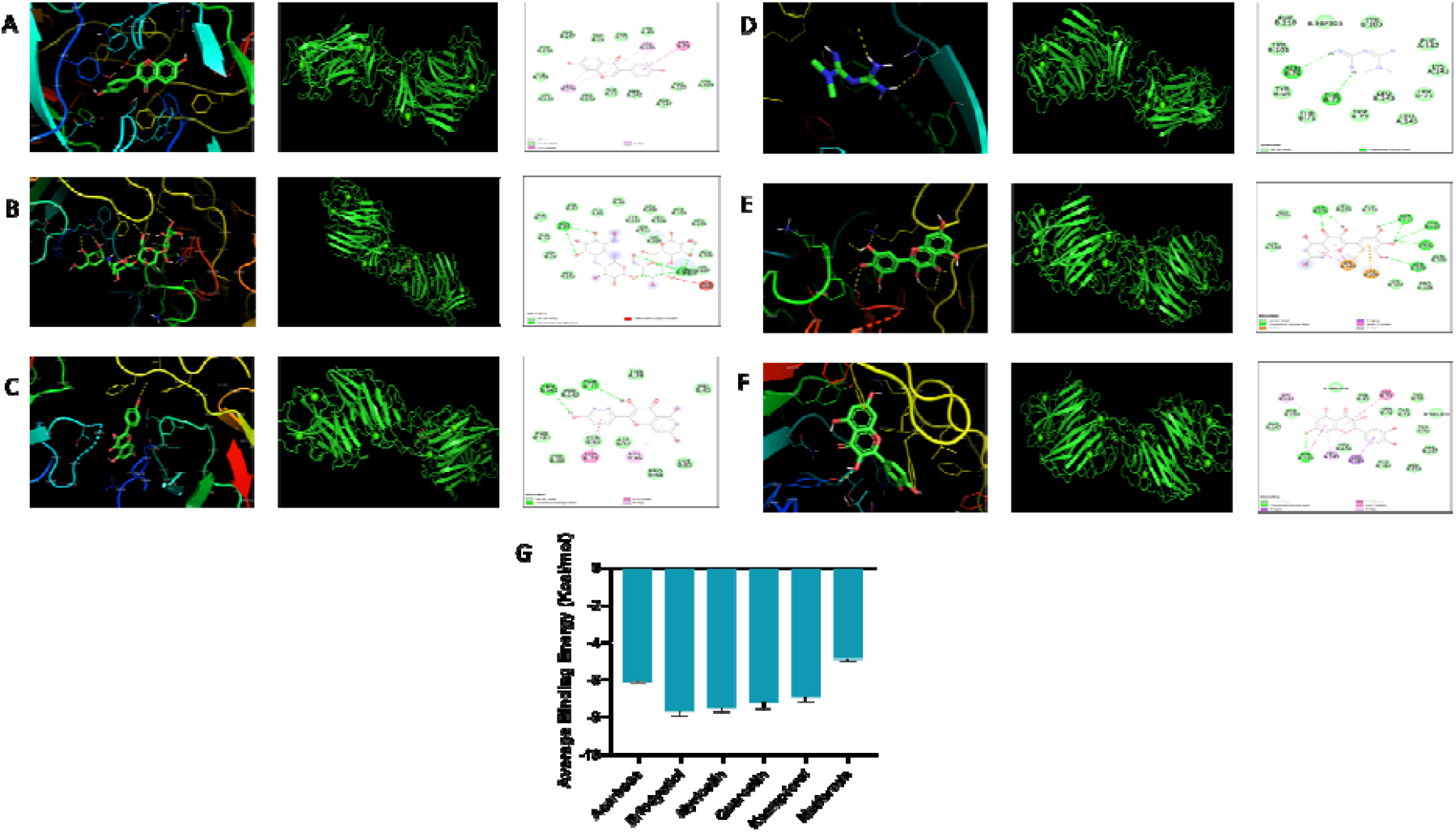
Computational visualization of the interaction of (A) Acarbose (B) Eriodyctiol (C) Myricetin (D) Quercetin (E) Kaempferol and (F) Metformin. (G) Binding affinity of selected ligands with α-amylase

### 3.4. Interactions of the ligands and Amylase

Eriodyctiol’s interaction with amylase is primarily characterized by hydrogen bonds with polar amino acid residues. Myricetin mainly forms hydrogen bonds with polar amino acids. In contrast, quercetin demonstrates both π-π stacking interactions with aromatic amino acid residues and hydrogen bonds with aspartic acid. The graphical representation of the protein-ligand interactions is illustrated in Figures 4A-H. Acarbose forms multiple types of interactions with the protein, including hydrogen bonds, van der Waals interactions, unfavorable donor-donor interactions, and π-alkyl interactions. Similarly, kaempferol exhibits several interactions, comprising π-π stacking interactions, hydrogen bonds, van der Waals interactions, and unfavorable donor-donor interactions. Lastly, metformin is associated with five hydrogen bonds, van der Waals interactions, and unfavorable donor-donor interactions.

### 3.5. Inhibition of PPA activity by flavonoids

Starch digestion by α-amylase is facilitated through the formation of a β-glycosyl enzyme intermediate, which is mediated by key acidic carboxylic residues within the enzyme’s active site, specifically Asp197, Glu233, and Asp300 ^34^. These residues, located in the catalytic cleft, play a critical role in the hydrolysis of starch. Polyphenolic compounds, particularly flavonoids, have been implicated as effective α-amylase inhibitors due to their interaction with the enzyme’s catalytic machinery^35, 36^. The inhibitory mechanism is largely driven by the presence of multiple hydroxyl groups on the B ring of the flavonoid skeleton, which facilitates hydrogen bonding with the enzyme’s active site residues^37^. Flavonoids such as myricetin and quercetin, belonging to the flavonol subgroup, exhibit strong inhibitory effects on α-amylase activity. Quercetin, with hydroxyl substitutions at positions 3, 5, 7 on the A ring and 3’, 4’ on the B ring, and myricetin, with additional hydroxyl substitution at position 5’, are found in various dietary sources, including onions, red wine, olive oil, berries, and grapes^37^. These hydroxyl groups allow for hydrogen bonding with the catalytic triad (Asp197, Glu233, and Asp300) within the enzyme’s active site, and the conjugated π-system of the flavonoid structure further stabilizes this interaction^35^. This mechanism underlies their inhibitory potential against α-amylase.

Eriodictyol, a flavanone, also exhibits inhibitory effects on α-amylase, though its structural differences from flavonols may influence its binding dynamics^38^.

All the tested flavonoids suppressed porcine pancreatic α-amylase (PPA) activity in a concentration-dependent manner. The IC_50_ values were determined to be 22 µM for eriodictyol, 19.96 µM for quercetin, and 24 µM for myricetin, as represented in Fig. 7 (A-C). These values demonstrate that quercetin is the most potent α-amylase inhibitor among the tested compounds, followed closely by myricetin and eriodictyol. The inhibitory activity of these flavonoids can be attributed to their structural composition, particularly the presence and positioning of hydroxyl groups. For instance, quercetin’s multiple hydroxyl groups allow for enhanced hydrogen bonding with the enzyme’s active site residues, contributing to its lower IC_50_ value.

**Figure 7:**
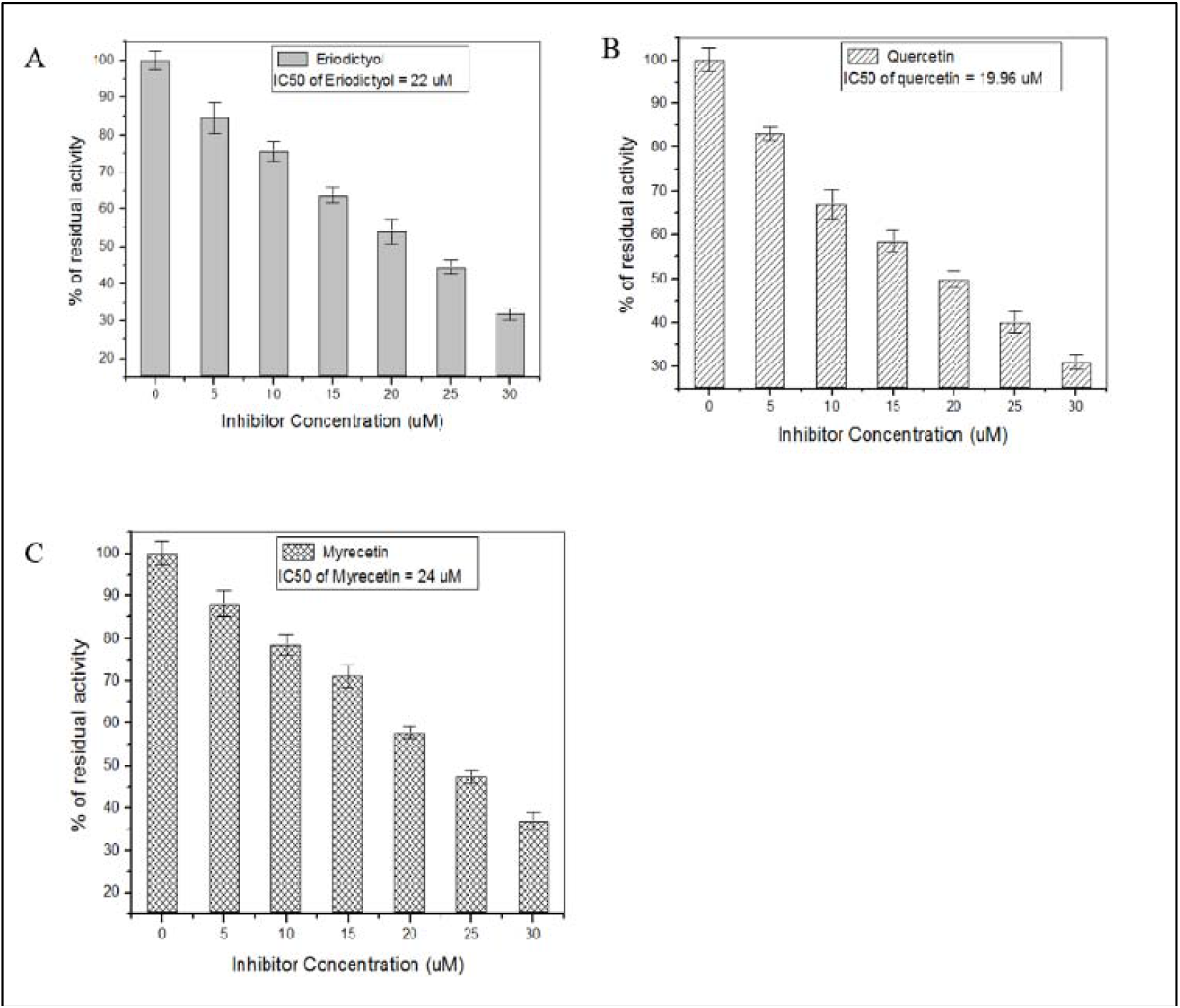
Inhibition of PPA activity by (A) Eriodyctiol (B) Quercetin and (C) Myricetin

**Figure 8:**
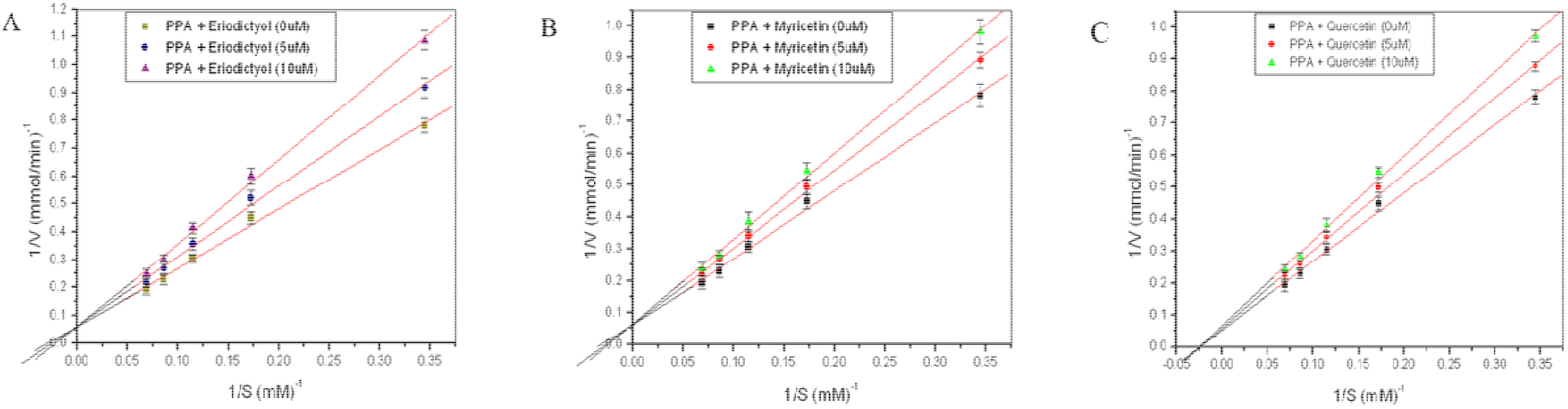
Lineweaver Burk plots of PPA inhibition by (A) Eriodyctiol (B) Myricetin and (C) Quercetin

### 3.6. Determination of the PPA inhibitory mode of action of flavonoids

To further elucidate the kinetics and the mode of inhibition of porcine pancreatic α-amylase (PPA) by the four flavonoids under study, a series of kinetic assays were conducted. Double reciprocal (Lineweaver-Burk) plots were generated to analyze enzyme kinetics both in the absence and presence of the flavonoids. The kinetic parameters, including the Michaelis-Menten constant (K_m_) and maximum velocity (V_max_), were derived from these plots and are summarized in Table 1.

**Table 1:** Kinetic parameters of PPA inhibition by flavonoids.

| Flavonoids | $K_M$ (mM) | Nature of inhibition |
| --- | --- | --- |
| Quercetin | 0.46 | Non- Competitive |
| Eriodictyol | 0.44, 0.42, 0.37 | Competitive |
| Myrecetin | 0.45, 0.41, 0.36 | Competitive |

The results indicate that eriodyctiol and myricetin act as competitive inhibitors of PPA. In competitive inhibition, the inhibitor binds directly to the active site of the enzyme, preventing substrate binding. This is reflected in the kinetic data where the Km value increases in the presence of the flavonoid, indicating a reduced affinity between PPA and its substrate. However, the V_max_ remains unchanged, which is characteristic of competitive inhibition. The competitive nature of eriodyctiol and myricetin suggests that these flavonoids interact directly with the enzyme’s active site, possibly through hydrogen bonding with critical residues such as Asp197, Glu233, and Asp300, as discussed previously. Their structural similarity, particularly the multiple hydroxyl groups in their flavonoid skeletons, enables these compounds to compete effectively with the substrate for active site binding. On the other hand, quercetin exhibits noncompetitive inhibition of PPA. This is evident from the kinetic data, where the V_max_ is reduced in the presence of quercetin, while the K_m_ remains relatively unchanged. The reduction in V_max_ signifies a decrease in the overall catalytic turnover, indicating that these flavonoids disrupt enzyme activity regardless of substrate concentration. This mode of inhibition implies that quercetin may interact with regions of the enzyme outside the catalytic site, inducing conformational changes that hinder the enzyme’s function. The differences in the inhibitory mechanisms between the flavonoids can be attributed to their structural variations. These findings are significant as they reveal different modes of inhibition exerted by structurally distinct flavonoids on PPA. Competitive inhibitors like eriodyctiol and myricetin could be more effective in situations where substrate concentrations are relatively low, as they directly compete with the substrate for binding. Noncompetitive inhibitors like quercetin, on the other hand, may provide a more consistent inhibitory effect regardless of substrate levels, making them potentially useful in maintaining steady enzyme inhibition under varying physiological conditions.

### 3.7. Effect of flavonoids on in vitro PPA inhibition in combination with acarbose

Acarbose belongs to the trestatin family with the acarviosine moiety. This pseudotetrasaccharide from *Actinoplanes* sp. is a polyhydroxylated aminocyclohexene derivative (valienamine) linked with a 6 deoxyglucose, bonded with a maltose moiety by α-1,4-linkage^39^. Being a competitive inhibitor of alpha-amylase, it has been routinely administered in the treatment of hyperglycemia. In acarbose, the valienamine moiety is attached to the binding subsite -1 and its strong inhibition is believed to be evolved from the binding of the valienamine group to the side chain of Asp197, Glu233, and Asp300^40^. Substitution of these polar amino acids may drop the catalytic activity of amylase indicating their importance in the binding. Keeping in mind the undesirable complications conceived of prolonged acarbose usage the need for an alternative therapeutic approach is never over. In the present study, % of PPA inhibition by two consecutive low concentrations of acarbose has been measured. In the presence of 3 µM and 6 µM acarbose, % of PPA inhibition is 9±1.2 and 16.2 ±1.3 % respectively. On the other hand, quercetin (10 µM) inhibits PPA by an amount of 24.3 % (Fig 9A). But when applied together with acarbose (3 µM and 6 µM), quercetin (10 µM) inhibits PPA by 33.6 ± .90 and 39 ± 1.1 % respectively. This stimulation of PPA inhibition by flavonoids may reduce the effective dose of acarbose synergistically. A similar *in vitro* report of the synergistic mode of action is also evident in the case of PPA inhibition by cyanidin-3-rutinoside ^41^. The present result is in agreement with the previous report indicating a common mechanistic approach is operative between PPA—acarbose complex and quercetin.

Amylase inhibition by acarbose occurs by binding acarbose to the substrate-binding site because of its structural similarity with starch and the kinetics have been characterized as competitive inhibition (data not shown). Among the four flavonoids, a similar type of binding characteristics has been observed for myricetin and eriodyctiol as manifested from their V_max_ and K_m_ values. In contrast, quercetin is a non-competitive inhibitor of PPA. Being a non-competitive inhibitor, they bind to a different site on PPA distinct from its substrate-binding site according to the following scheme with some conformational change of PPA leading to the diminished substrate binding.

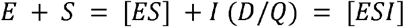

Or

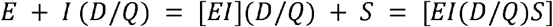

So, in the presence of quercetin, binding can be accompanied by acarbose at the same time during the reaction with an outcome of magnified PPA inhibition. This eventually reduces the dose of acarbose for *in vitro* PPA inhibition in combination with flavonoids which are noncompetitive inhibitors of PPA. The effect can be of clinical significance as it may ameliorate the undesirable side effects of acarbose used at a high dose for the treatment of diabetic patients.

### 3.8. Determination of thermodynamic parameters of Quercetin binding to PPA

Fig. 10 (A) shows a clear downward trend in fluorescence, which can be attributed to quenching. The increasing fluorescence quenching with higher Quercetin concentrations suggests that the binding of Quercetin to PPA intensifies as its concentration rises, likely due to direct interaction with the fluorophore within PPA. Alternatively, the quenching could result from a conformational change in PPA induced by Quercetin, causing the protein to fold and bury its fluorescent residues, leading to reduced fluorescence. To clarify the underlying mechanism, the thermodynamic parameters of the interaction were analyzed as represented in Fig. 10 (B). Through an isothermal titration colorimetry assay, we obtained the parameters as

- N (stoichiometry): 0.331 ± 0.114, indicating less than 1 ligand molecule binds per protein.
- K (association constant): 2.515E5 ± 1.38E5 M□^1^, indicating moderate to strong binding affinity.
- ΔH (enthalpy change): -1345 ± 553.1 kcal/mol, which is exothermic, suggesting a favorable enthalpy contribution to binding.
- ΔS (entropy change): 20.3 cal/mol·K, indicating a small entropy gain, possibly due to desolvation effects or structural flexibility.

The data suggests a single binding event with strong affinity (K = 2.515E5 M□^1^), driven primarily by enthalpic interactions (ΔH = -1345 kcal/mol), with minimal entropy contribution (ΔS = 20.3 cal/mol·K). The stoichiometric interaction (N = 0.331) suggests a sub-stoichiometric interaction, possibly indicating that less than one full ligand molecule binds per macromolecule, or there is cooperative or complex binding.

**Figure 10:**
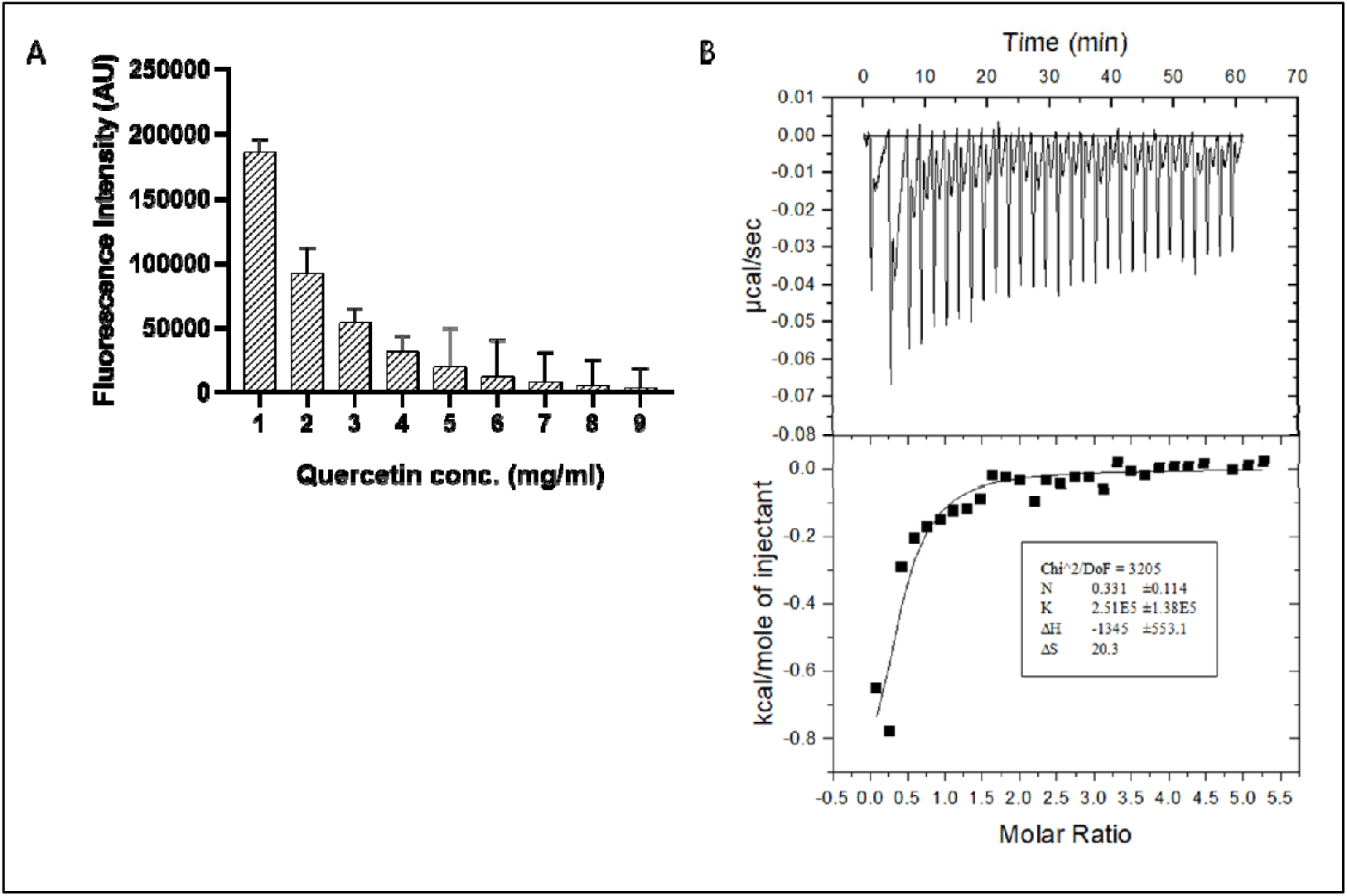
Binding interactions of PPA and Quercetin through (A) fluorescence quenching and (B) isothermal titration colorimetry assay

### 3.9. In vitro anti-lipid peroxidation assay in the presence of flavonoids

Anti-lipid peroxidation activity is very pronounced and a key event for study in this connection. Oxidative stress being a common factor for several diseases like cancer, diabetes, ageing, inflammatory conditions, hepatic/neural disorders, and atherosclerosis generates several free rad A like nitric oxide (NO), and superoxide radicals (O_2_^.-^), and hydroxyl radical (OH.). These reactive oxygen species (ROS) formation is stimulated in the presence of free metal ions damaging cell membranes and inactivating enzymes ^42^. Flavonoids with lower redox potentials reduce harmful free radicals by donation of H atom and can chelate metal ions to attenuate free radical generation ^43^. Quercetin, epicatechin, and rutin in this regard need special mention with their 3^/^,4^/^ catechol structure in the B ring requisite for enhanced metal chelating ability accounting for anti-lipid peroxidation profile ^44^. The resulting fairly stable orthosemiquinone radical from the flavonoids can act as strong scavengers offering an impasse to lipid peroxidation ^45^. To eliminate any possible interference by acarbose on the inhibitory activity of flavonoids during lipid peroxidation, quercetin was tested in the absence and presence of acarbose. Lipid peroxidation assay has been performed in the presence of Quercetin at its IC_50_ values (26 and 20 µM respectively) for PPA inhibition. No remarkable reduction in the inhibition of lipid peroxidation by Quercetin has been manifested when carried out in the presence of 18 µM acarbose (1/2 of the IC_50_ value of acarbose for PPA inhibition). The extent of lipid peroxidation induced by FeSO_4_ has been taken as 100% which has been diminished to 81.5% in the presence of 20 µM Quercetin (IC_50_ for PPA inhibition).Therefore, there is a reduction of lipid peroxidation in the presence of Quercetin which is not interrupted by the presence of 18 µM acarbose used for *in vitro* combination study during PPA inhibition along with 20 µM Quercetin as shown in Fig. 11 (A).

**Figure 11:**
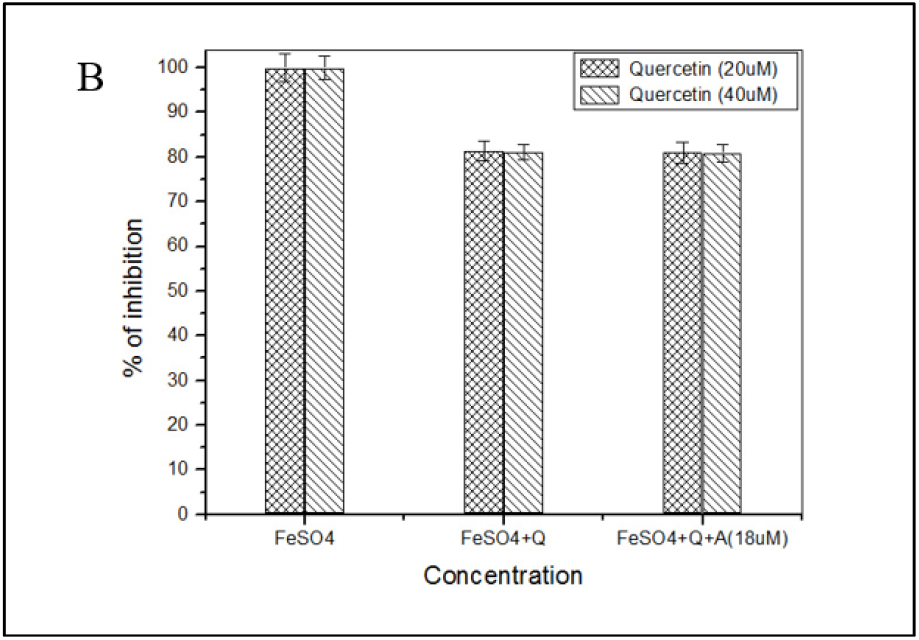
Inhibition of lipid peroxidation by (A) Quercetin with and without Acarbose

**Figure 12:**
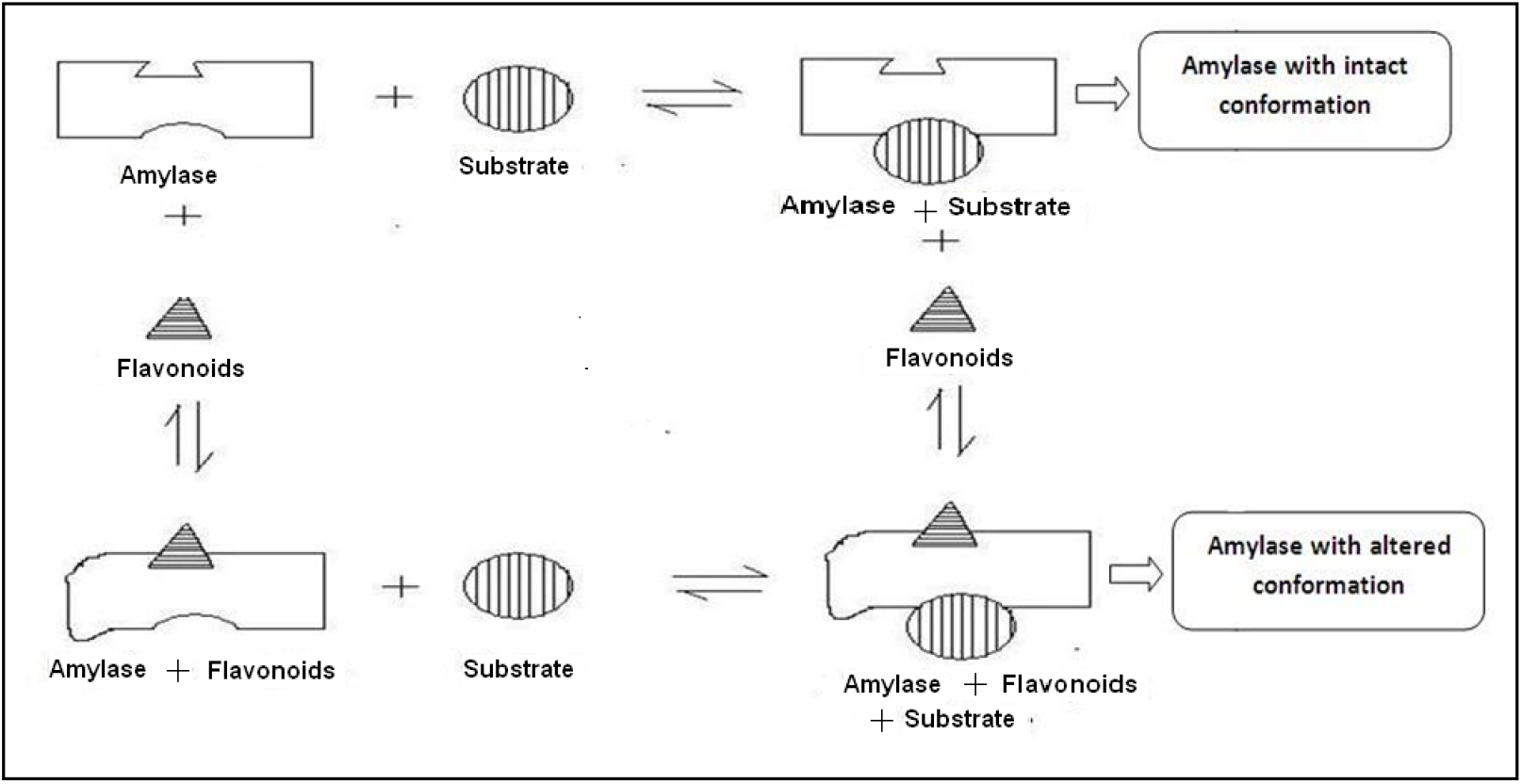
Schematic representation of enhanced inhibition of PPA by acarbose in the presence of flavonoids

The apparent difference in the extent of inhibition of lipid peroxidation by these two flavonoids is due to the difference in their concentrations (26 vs 20 µM) used for the assay. The result is indicative in support of the simultaneous use of acarbose and flavonoids during PPA inhibition as the lipid peroxidation potential of Quercetin has been protected in the presence of acarbose. Doubling the dose of acarbose (IC_50_ = 36 µM) does not make any significant difference in the extent of lipid peroxidation by D and Q (data not shown). Based on the above result a model has been proposed to elucidate the mechanistic pathway of PPA inhibition by acarbose in the presence of quercetin (Fig.12).

## 4. Conclusion

This study highlights the potent in vitro inhibitory effects of various flavonoids on porcine pancreatic α-amylase (PPA), which can potentially be leveraged in the management of diabetes. The findings suggest that flavonoids could be used synergistically with existing amylase inhibitors like acarbose, allowing for a reduction in the required dose of acarbose, thereby minimizing its associated side effects while maintaining therapeutic efficacy. Given the extensive health-promoting properties of flavonoids, including antimicrobial, anticancer, antidiabetic, and anti-lipid peroxidation activities, their regular inclusion in the human diet could have far-reaching benefits. Flavonoids occupy a crucial position in the management of several critical diseases, including diabetes and cancer, with minimal side effects as supported by numerous earlier reports. However, while these *in vitro* findings provide valuable insights into the mechanism of PPA inhibition by flavonoids, further *in vivo* studies are essential to confirm their therapeutic potential in humans. Future research should focus on evaluating the efficacy of flavonoids in diabetic patients, particularly in combination with acarbose or other amylase inhibitors, to determine their impact on postprandial hyperglycemia and overall glucose management. Such investigations, involving diabetic volunteers, will be crucial in translating these *in vitro* observations into clinical practice, where flavonoid-based therapies could play a pivotal role in improving glycemic control and reducing the burden of diabetes.

## Abbreviations

(E): Eriodyctiol
(M): Myricetin
(Q): Quercetin
(PPA): porcine pancreatic amylase

## Acknowledgment

The author thanks the Department of Biotechnology and M.Tech student, Mr. Sujan Maity for his support and work.

## References

(1) Cuschieri, S. Type 2 diabetes–An unresolved disease across centuries contributing to a public health emergency. Diabetes & Metabolic Syndrome: Clinical Research & Reviews 2019, 13 (1), 450–453.

(2) Abdul Basith Khan, M.; Hashim, M. J.; King, J. K.; Govender, R. D.; Mustafa, H.; Al Kaabi, J. Epidemiology of type 2 diabetes—global burden of disease and forecasted trends. Journal of epidemiology and global health 2020, 10 (1), 107–111.

(3) Tundis, R.; Loizzo, M. R.; Menichini, F. Natural products as α-amylase and α-glucosidase inhibitors and their hypoglycaemic potential in the treatment of diabetes: an update. Mini Rev. Med. Chem. 2010, 10 (4), 315–331.

(4) Galicia-Garcia, U.; Benito-Vicente, A.; Jebari, S.; Larrea-Sebal, A.; Siddiqi, H.; Uribe, K. B.; Ostolaza, H.; Martín, C. Pathophysiology of type 2 diabetes mellitus. International journal of molecular sciences 2020, 21 (17), 6275.

(5) Mansour, A.; Mousa, M.; Abdelmannan, D.; Tay, G.; Hassoun, A.; Alsafar, H. Microvascular and macrovascular complications of type 2 diabetes mellitus: Exome wide association analyses. Frontiers in Endocrinology 2023, 14, 1143067.

(6) Bhatti, J. S.; Sehrawat, A.; Mishra, J.; Sidhu, I. S.; Navik, U.; Khullar, N.; Kumar, S.; Bhatti, G. K.; Reddy, P. H. Oxidative stress in the pathophysiology of type 2 diabetes and related complications: Current therapeutics strategies and future perspectives. Free Radical Biology and Medicine 2022, 184, 114–134.

(7) Sales, P. M.; Souza, P. M.; Simeoni, L. A.; Magalhães, P. O.; Silveira, D. α-Amylase inhibitors: a review of raw material and isolated compounds from plant source. J. Pharm. Pharm. Sci. 2012, 15 (1), 141–183.

(8) Inzucchi, S. E. Oral antihyperglycemic therapy for type 2 diabetes: scientific review. Jama 2002, 287 (3), 360–372.

(9) Chakrabarti, R.; Rajagopalan, R. Diabetes and insulin resistance associated disorders: disease and the therapy. Curr. Sci. 2002, 1533–1538.

(10) Phan, M. A. T.; Wang, J.; Tang, J.; Lee, Y. Z.; Ng, K. Evaluation of α-glucosidase inhibition potential of some flavonoids from Epimedium brevicornum. LWT 2013, 53 (2), 492–498.

(11) Hung, H. Y.; Qian, K.; Morris-Natschke, S. L.; Hsu, C. S.; Lee, K. H. Recent discovery of plant-derived anti-diabetic natural products. Nat. Prod. Rep. 2012, 29 (5), 580–606.

(12) Hedrington, M. S.; Davis, S. N. Considerations when using alpha-glucosidase inhibitors in the treatment of type 2 diabetes. Expert opinion on pharmacotherapy 2019, 20 (18), 2229–2235.

(13) Lankisch, M.; Layer, P.; Rizza, R. A.; DiMagno, E. P. Acute postprandial gastrointestinal and metabolic effects of wheat amylase inhibitor (WAI) in normal, obese, and diabetic humans. Pancreas 1998, 17 (2), 176–181.

(14) Boivin, M.; Flourie, B.; Rizza, R. A.; Go, V. L. W.; DiMagno, E. P. Gastrointestinal and metabolic effects of amylase inhibition in diabetics. Gastroenterology 1988, 94 (2), 387–394.

(15) Eyre, H.; Kahn, R.; Robertson, R. M.; Clark, N. G.; Doyle, C.; Hong, Y.; Gansler, T.; Glynn, T. Preventing cancer, cardiovascular disease, and diabetes: a common agenda for the American Cancer Society, the American Diabetes Association, and the American Heart Association. Circulation 2004, 109 (25), 3244–3255.

(16) Formica, J. V.; Regelson, W. Review of the biology of quercetin and related bioflavonoids. Food Chem. Toxicol. 1995, 33 (12), 1061–1080.

(17) Di Magno, L.; Di Pastena, F.; Bordone, R.; Coni, S.; Canettieri, G. The mechanism of action of biguanides: New answers to a complex question. Cancers 2022, 14 (13), 3220.

(18) Bolen, S.; Feldman, L.; Vassy, J.; Wilson, L.; Yeh, H. C.; Marinopoulos, S.; Wiley, C.; Selvin, E.; Wilson, R.; Bass, E. B. Systematic review: comparative effectiveness and safety of oral medications for type 2 diabetes mellitus. Ann. Intern. Med. 2007, 147 (6), 386–399. DOI: 10.7326/0003-4819-147-6-200709180-00178.

(19) Andrès, E.; Noel, E.; Goichot, B. Metformin-associated vitamin B12 deficiency. Arch. Intern. Med. 2002, 162 (19), 2251–2252.

(20) Pérez-Ros, P.; Navarro-Flores, E.; Julián-Rochina, I.; Martínez-Arnau, F. M.; Cauli, O. Changes in Salivary Amylase and Glucose in Diabetes: A Scoping Review. Diagnostics 2021, 11 (3), 453. DOI: 10.3390/diagnostics11030453 PubMed.

(21) Malla, A.; Gupta, S.; Sur, R. Inhibition of lactate dehydrogenase A by diclofenac sodium induces apoptosis in H e L a cells through activation of AMPK. The FEBS Journal 2024.

(22) Ainscow, E. K.; Zhao, C.; Rutter, G. A. Acute overexpression of lactate dehydrogenase-A perturbs beta-cell mitochondrial metabolism and insulin secretion. Diabetes 2000, 49 (7), 1149–1155.

(23) Avogaro, A.; Toffolo, G.; Miola, M.; Valerio, A.; Tiengo, A.; Cobelli, C.; Del Prato, S. Intracellular lactate-and pyruvate-interconversion rates are increased in muscle tissue of non-insulin-dependent diabetic individuals. J. Clin. Investig. 1996, 98 (1), 108–115. DOI: 10.1172/JCI118754.

(24) Burlingham, B. T.; Widlanski, T. S. An intuitive look at the relationship of Ki and IC50: a more general use for the Dixon plot. J. Chem. Educ. 2003, 80 (2), 214.

(25) Baskar, R.; Rajeswari, V.; Kumar, T. S. In vitro antioxidant studies in leaves of Annona species. NOPR 2007.

(26) Brayer, G. D.; Sidhu, G.; Maurus, R.; Rydberg, E. H.; Braun, C.; Wang, Y.; Nguyen, N. T.; Overall, C. M.; Withers, S. G. Subsite mapping of the human pancreatic α-amylase active site through structural, kinetic, and mutagenesis techniques. Biochemistry 2000, 39 (16), 4778–4791.

(27) Trott, O.; Olson, A. J. AutoDock Vina: improving the speed and accuracy of docking with a new scoring function, efficient optimization, and multithreading. J. Comput. 2010, 31 (2), 455–461.

(28) Schrödinger, L. The PyMol Molecular Graphics System, Versión 1.8. Thomas Hold. 2015.

(29) Biovia, D. S. B. Discovery Studio Modelling Environment, Release 4.5, Accelrys Softw. Inc: 2015.

(30) Lo Piparo, E.; Scheib, H.; Frei, N.; Williamson, G.; Grigorov, M.; Chou, C. J. Flavonoids for controlling starch digestion: structural requirements for inhibiting human α-amylase. J. Med. Chem. 2008, 51 (12), 3555–3561.

(31) Kim, J. S.; Kwon, C. S.; Son, K. H. Inhibition of alpha-glucosidase and amylase by luteolin, a flavonoid. Biosci. Biotechnol. Biochem. 2000, 64 (11), 2458–2461.

(32) Graham, H. N. Green tea composition, consumption, and polyphenol chemistry. Preventive medicine 1992, 21 (3), 334–350.

(33) Chatterjee, P.; Chandra, S.; Dey, P.; Bhattacharya, S. Evaluation of anti-inflammatory effects of green tea and black tea: A comparative: in vitro: study. Journal of advanced pharmaceutical technology & research 2012, 3 (2), 136–138.

(34) Neves, R. P.; Fernandes, P. A.; Ramos, M. J. Role of enzyme and active site conformational dynamics in the catalysis by α-amylase explored with QM/MM molecular dynamics. Journal of Chemical Information and Modeling 2022, 62 (15), 3638–3650.

(35) Akkarachiyasit, S.; Yibchok-Anun, S.; Wacharasindhu, S.; Adisakwattana, S. In vitro inhibitory effects of cyandin-3-rutinoside on pancreatic α-amylase and its combined effect with acarbose. Molecules 2011, 16 (3), 2075–2083.

(36) Van Acker, S. A. B. E.; Tromp, M. N. J. L.; Griffioen, D. H.; Van Bennekom, W. P.; Van Der Vijgh, W. J. F.; Bast, A. Structural aspects of antioxidant activity of flavonoids. Free Radic. Biol. Med. 1996, 20 (3), 331–342.

(37) Kumar, S.; Pandey, A. K. Chemistry and biological activities of flavonoids: an overview. The scientific world journal 2013, 2013 (1), 162750.

(38) Ku, Y.-S.; Ng, M.-S.; Cheng, S.-S.; Lo, A. W.-Y.; Xiao, Z.; Shin, T.-S.; Chung, G.; Lam, H.-M. Understanding the composition, biosynthesis, accumulation and transport of flavonoids in crops for the promotion of crops as healthy sources of flavonoids for human consumption. Nutrients 2020, 12 (6), 1717.

(39) Tsunoda, T.; Samadi, A.; Burade, S.; Mahmud, T. Complete biosynthetic pathway to the antidiabetic drug acarbose. Nature Communications 2022, 13 (1), 3455.

(40) Yoon, S.-H.; Robyt, J. F. Study of the inhibition of four alpha amylases by acarbose and its 4IV-α-maltohexaosyl and 4IV-α-maltododecaosyl analogues. Carbohydrate research 2003, 338 (19), 1969–1980.

(41) Speciale, A.; Saija, A.; Bashllari, R.; Molonia, M. S.; Muscarà, C.; Occhiuto, C.; Cimino, F.; Cristani, M. Anthocyanins as modulators of cell redox-dependent pathways in non-communicable diseases. Curr. Med. Chem. 2020, 27 (12), 1955–1996.

(42) ALrefai, A. A.; Alsalamony, A. M.; Fatani, S. H.; Kamel, H. F. Effect of variable antidiabetic treatments strategy on oxidative stress markers in obese patients with T2DM. Diabetology & Metabolic Syndrome 2017, 9, 1–8.

(43) Mishra, A.; Tyagi, C.; Pandey, B.; Chakraborty, O.; Kumar, A.; Jain, A. Structural insights into the mode of action of plant flavonoids as anti-oxidants using regression analysis. Proceedings of the National Academy of Sciences, India Section B: Biological Sciences 2016, 86, 1023–1036.

(44) Dajas, F.; Arredondo, F.; Echeverry, C.; Ferreira, M.; Morquio, A.; Rivera, F. J. C. N. Flavonoids and the brain: Evidences and putative mechanisms for a protective capacity. Curr. Neuropharmacol. 2005, 3 (3), 193–205.

(45) Fauconneau, B.; Waffo-Teguo, P.; Huguet, F.; Barrier, L.; Decendit, A.; Merillon, J.-M. Comparative study of radical scavenger and antioxidant properties of phenolic compounds from Vitis vinifera cell cultures using in vitro tests. Life sciences 1997, 61 (21), 2103–2110.

